# Automated speech detection in eco-acoustic data enables privacy protection and human disturbance quantification

**DOI:** 10.1101/2022.02.08.479660

**Authors:** Benjamin Cretois, Carolyn Rosten, Sarab S. Sethi

## Abstract

Eco-acoustic monitoring is increasingly being used to map biodiversity across huge scales, yet little thought is given to the privacy concerns and potential scientific value of inadvertently recorded human speech. Automated speech detection is possible using Voice Activity Detection (VAD) models, but existing approaches have been developed for indoor or urban use cases, rather than diverse natural soundscapes. In this study we used a data augmentation approach to create ecoVAD, a convolutional neural network designed for robust voice detection in eco-acoustic data. We performed playback experiments using speech samples from a woman, man, and child in two ecosystems in Børsa, Norway, and showed that ecoVAD was able to accurately detect voices at distances of up to 20m – at which point the speech was unintelligible. We compared ecoVAD with two existing VAD models and found that ecoVAD consistently outperformed the state-of-the-art (mean F1 scores: ecoVAD, 0.917; pyannote, 0.890; WebRTC VAD, 0.876). Using long-term passive recordings from a popular hiking location in Bymarka, Norway, we found that the frequency of speech detections was linked closely to peak traffic hours (using bus timings) demonstrating how VAD models can be used to quantify human activity with a fine temporal resolution. Anonymising audio data effectively using VAD models will allow eco-acoustic monitoring to continue to deliver invaluable ecological insight at scale, whilst minimising the risk of data misuse. Furthermore, using speech detections as a fine scale measure of human disturbance opens new possibilities for studying subtle human-wildlife interactions on the vast scales made possible by eco-acoustic monitoring technology.

**Author Summary:** Eco-acoustic monitoring (i.e., recording and analysing the sounds of an ecosystem) allows us to monitor biodiversity and ecological health on vast spatial and temporal scales. However, as networks of audio recorders are increasingly being deployed across the world by scientists and land managers, little thought has been given to the implications of inadvertently recorded human speech in these datasets. We developed a neural-network model to detect and remove speech from audio data, and showed it performed more accurately than existing state-of-the-art models when evaluated on eco-acoustic datasets. Furthermore, we showed frequency of voice detections could be used to monitor human pressure on an ecosystem. Our work paves the way for ethical and responsible processing of eco-acoustic data and provides a novel approach to studying human wildlife interactions on a large scale with high temporal resolution.

## Introduction

Land-use change and global warming are impacting the natural world at an ever-increasing rate (1,2). By exploiting advances in sensor technology and data analysis techniques, scientists are now able to monitor and understand the resulting ecological changes both in detail and at unprecedented scales (3). Eco-acoustic monitoring is a particularly promising monitoring approach which offers high resolution data across a wide range of taxa over long time periods (4,5). However, whilst the field of eco-acoustic monitoring is expanding rapidly, surprisingly little attention has been paid to an almost universal issue: the impact and implications of inadvertently recorded human speech.

Eco-acoustic data is collected easily and inexpensively and can be analysed with increasingly sophisticated analytical techniques (6–9). The maturity of the technology has led to autonomous monitoring networks being deployed across vast landscapes, and even across full nations (10,11). Whilst the primary focus is to derive ecological data such as species distributions or community assemblages, some recording of human speech is inevitable. The presence of identifiable human speech raises serious ethical questions regarding data privacy and opens the door to potentially nefarious uses of eco-acoustic data; yet it is has not to date been discussed in the literature. Manual filtering of data or obtaining prior consent is not appropriate nor practically feasible in most cases, and therefore an automated approach to anonymising eco-acoustic data is required.

Acoustic data can be anonymised by blurring or removing sections of audio containing identifiable speech. Solutions to the most challenging step, Voice Activity Detection (VAD), have been explored in other fields of acoustic data analysis (12,13). However, applications tend to be either in situations where the speaker is close to or speaking directly into a microphone (14), or in urban environments with relatively low levels of homogenous background noise (15). Detecting speech in highly diverse and noisy passively collected eco-acoustic data is a more difficult task; especially since common components such as bird calls can overlap in frequency with speech and may trigger false positives (16). In-depth evaluation and adaptation of existing state-of-the-art VAD models is required to ensure anonymisation can be done reliably given the uniquely challenging nature of eco-acoustic data.

Recorded human speech, whilst undesirable from a privacy standpoint, has the potential to provide invaluable data on levels of human disturbance. Eco-acoustic data has previously been used to estimate levels of human disturbance at a coarse soundscape level (17,18). Nonetheless, voice activity detection would provide as a by-product far more detailed information on exactly when and where humans were present, as well as presenting the opportunity to estimate human population sizes and demographics (19,20). Quantifying human disturbance at this resolution and level of detail may shed light on subtle human-wildlife interactions that would otherwise be hidden when using coarser measures of disturbance.

In this study we present a novel approach to voice activity detection which is tailor-made for anonymisation of eco-acoustic data. We trained Convolutional Neural Networks (CNN) on synthetic datasets comprised of human voices augmented with typical background sounds encountered in eco-acoustic data. We measured the ability of our model to anonymise real data with controlled field experiments and evaluated its ability to generalise across different landscapes. Finally, we explored the potential for VAD models to quantify human disturbance in a Norwegian landscape regularly used for recreational hiking purposes. Results were validated by comparing the frequency of voice detections with bus arrival and departure times – used here as an indicator for true human activity.

## 2. Methods

### 2.1. Playback experiment dataset

We selected two sites located in Børsa, Norway to collect training, validation, and test audio data for playback experiments. Each site represented a different landscape type, namely a forest and a semi-natural grassland. At each site we deployed an Audiomoth version 1.1.1 device (7) that recorded audio data for five consecutive days during July 2021. We placed the Audiomoths at least 100m from the closest hiking trail to avoid unnaturally high levels of human noise pollution. Audio was recorded at a sampling frequency of 44.1kHz continuously in chunks of 55 seconds and files were saved in the WAV format. Four days (N=5700 soundscape audio files) were used for training and one day (N=1440 soundscape audio files) was retained for validation.

To test the VAD models, we recorded speech from three different speakers (a female, a male, and a child) in a soundproof environment. Each person read the exact same sentence at a volume used during a normal conversation (Supplementary C) resulting in a recording of approximatively 30 seconds. We played the recorded speech using a JBL Xtreme 2 portable speaker, with the volume calibrated such that the male voice registered at 60dB SPL (i.e., the volume of real speech).

Playbacks were performed at distances of 1, 5, 10, and 20 meters from an Audiomoth device in both the forest and semi-natural grasslands. Samples of speech from each of the three speakers were played at each distance, both facing and from behind the recording device. In total, we collected 48 two-minute recordings, each containing approximatively 30 seconds of speech and 1 min 30 s of ambient soundscapes. The recorded audio was divided into three second clips and each was manually labelled as “speech” or “no speech”.

### 2.2. Soundscape data for human disturbance quantification

To measure the performance of our model in a passive monitoring scenario we collected data from the Bymarka forest near Trondheim, Norway. This forest is particularly popular for weekend hikes and outdoors activities, so we expected significant human activity and speech. Two audiomoth version 1.1.1 devices were deployed for five days each and data was recorded in files of 55 seconds and at a sampling frequency of 44.1 kHz. The first audiomoth (Forest 1) was located closer to a road (distance from the nearest road = 40 meters) while the second audiomoth (Forest 2) was located further into the forest (distance from the nearest road = 140 meters).

Five days were used for model training, and we used one day as a validation dataset (N=6140, 1440 soundscape audio files for training and validation respectively). We retained the Saturday 2^nd^ and Sunday 3^rd^ of October 2021 for both audiomoths as test data (i.e. four days in total; N=5700 soundscape audio files).

### 2.3 Data augmentation pipeline

Ecosystems have soundscapes which are far more diverse than the environments in which VAD models are typically deployed. Common sounds come from both biotic (e.g., species vocalizing or moving) and abiotic (e.g., rain, wind, motors) sources – each of which can vary from site to site and can confound sound event detection models. Existing human speech datasets used to train VAD models, however, do not capture this diversity of background sounds. Therefore, we took a data augmentation approach and combined several datasets to train a custom CNN based VAD model on a large synthetic dataset which is representative of typical eco-acoustic data.

To generate the augmented dataset, we divided the 55 seconds soundscape recordings into non-overlapping three seconds segments. Each three second file was processed in a pipeline that added human voices (Librispeech, 21), and background noises: animal vocalizations (BirdClef2017, 22), and environmental and anthropogenic sounds (ESC50, 23) (Supplementary A). We added both human speech and background noises to 25%, human speech only to 25%, and background noises only to 45% of the three second records. We did not add any human speech or background noises to 5% of the records (Figure 1a).

**Figure 1.**
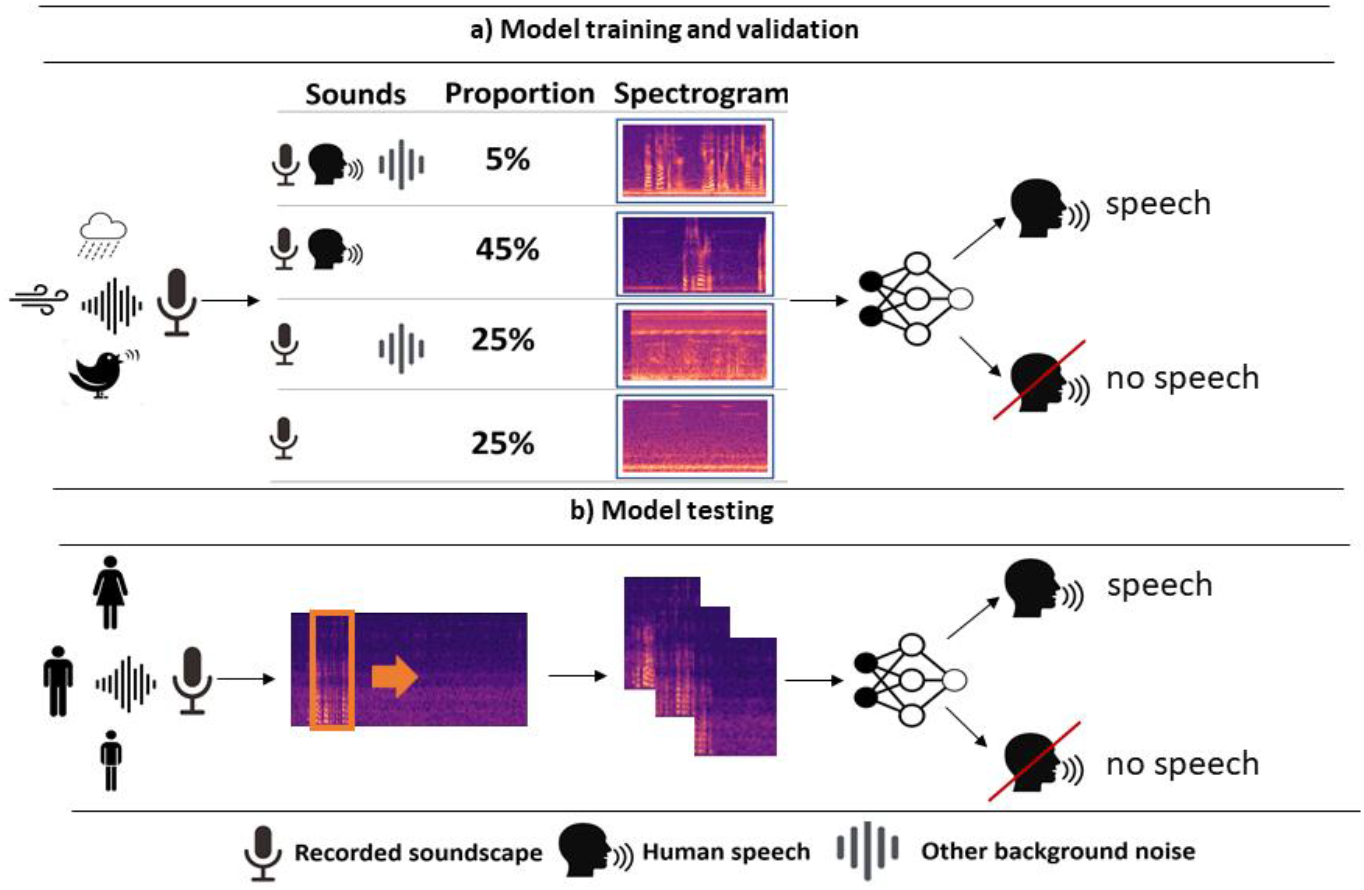
Schematic overview of our approach to automated human speech detection in eco-acoustic data. **a)** Training and validation data was created using a data augmentation approach, where existing datasets of human speech and typical background noises were overlaid with varying probabilities on baseline ecosystem soundscape data. A convolutional neural network was trained to detect human speech using this augmented data. **b)** We performed playback experiments using recorded speech samples from a man, woman, and child to measure the accuracy of our model. We also evaluated the performance of our model on longer passively recorded eco-acoustic data.

When only one sound type was added (speech or background noise) waveforms were scaled by a parameter, α, such that the amplitude of the added audio varied randomly in the range [-56.16, -8.3] dBFS (Eq. 1, Supplementary B). If background noise and human speech were both added to a 3 second audio file, a parameter *β* drawn from a uniform distribution in the range [0.1, 0.9] was additionally used (Eq. 2). We also added a fade in and fade out effect of 500ms for the human speech clip to simulate a human speaker moving towards and past the recording device.

We randomly subsampled three seconds of audio from each human speech clip so that the clip could begin in the middle of an utterance. To simulate speech being partially captured, we also included a probability of random shifts in start time for both the human speech and background noises. Thus, speech or background noise clips were started anytime between [0, 2 seconds] in the three seconds segment for a duration of at least one second.

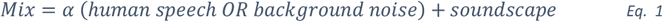

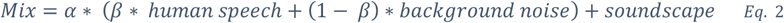

We tested the efficacy of data augmentation in eco-acoustic VAD models was by training and validating models on a) a fully augmented dataset (ecoVAD), b) a dataset containing only the natural soundscape (Soundscape aug), c) a dataset containing only background noises (Noise aug), and d) a dataset that had not been augmented (No aug).

To generate Soundscape aug we removed all background noises from the pipeline, the dataset thus containing 50% of segments with human speech and soundscape background with varying SNR (Eq. 1) and 50% of segments with unmodified soundscape. Noise aug was generated by replacing the soundscape background by white noise whose volume was fixed at -50 dBFS, the root mean square (RMS) volume of the raw recorded soundscapes. The rest of the pipeline was unchanged. Finally, No aug was generated by removing all background noises and replacing the soundscape by white noise as described above. We made sure that the same number of training and validation files was used for all augmentation types and that, where appropriate, each augmentation type included the same speech and background noise files.

### 2.4 CNN model architecture and training

All custom CNN models were based on the VGG11 architecture (24). We changed the number of input neurons so that the model accepted fixed-size 128 × 96 images containing a single colour channel. To reduce overfitting, we also added batch normalization after each convolutional layer and a dropout of 0.5 for the fully connected layers.

We converted each three augmented second segment into a Mel spectrogram that was used as an input to the model. Acknowledging that our binary classification task did not require very high temporal resolution, we selected a Fast Fourier Transform (FFT) window size of 64 ms (1024 samples at 16 kHz) and an overlap of 50% (hop length of 512). We performed frequency compression using a Mel scale with 96 bands. Finally, we normalized the Mel spectrograms along each frequency bin as this has been shown to significantly improve performance for audio classification tasks (25). These steps resulted in a Mel spectrogram of dimension 96 × 128 pixels that was used as an input for the model (Figure 1a).

Because the playback dataset contained two different soundscapes (Forest and Semi-natural grassland) we trained one model for each data augmentation type (i.e. ecoVAD, Noise aug, Soundscape aug and No aug) and for each soundscape type. We also trained models using data from both soundscape types put together (Forest + Semi-natural grassland). This resulted in a total of 12 models for the playback dataset and four models for the Bymarka dataset.

All models were trained using a learning rate of 0.1, 0.01 and 0.001 with a decay of 0.1 every 20 epochs. An early stopping strategy based on the validation loss was used to stop the training of the model if it was complete before the maximum of 200 epochs. Performance of the models was evaluated on the validation dataset, and we used the Area Under the Receiver Operating Characteristic Curve (ROC AUC, herein AUC) to select the best performing learning rates per data augmentation type and landscape type. The performances of the best models were then evaluated on the test datasets.

### 2.5 CNN model testing

Model predictions on the test data were done using a sliding window with an overlap of two seconds between an input audio segment and the previous one, resulting in one prediction per second of audio (Figure 1b). Since we had ground truth labels for the playback dataset, we measured accuracy using AUC.

Similar ground truth labels were unavailable for the Bymarka dataset as the length of the recording period made manual labelling infeasible. To estimate model performance for each augmentation type, we randomly selected 100 detections per audiomoth that we manually listened to confirm whether the detections were correct (i.e., contained any human speech). Comparing the number of true positive and false negative yielded the precision for each model.

We also counted the number of detections the model made per hour. The number of detections was normalized using Min-Max scaling to produce a scaled measure of relative human pressure across the test period. Finally, we compared the number of detections of the models with the arrival and departure of busses at the nearest bus stop from the audiomoth (distance to nearest bus stop = 135, 235 meters for audiomoth Forest 1 and 2 respectively). The bus timetable was obtained from the bus company website (https://www.atb.no/en/)

### 2.6. Comparison with other VAD models

We compared the performance of our model with two existing state-of-the-art VAD models. Based on their widespread usage and differing approaches, we used pyannote v1.1.1 (26), a CNN based on a PyanNet architecture and Google WebRTC VAD v2.0.10 (27), a machine learning algorithm based on a Gaussian Mixture Model. Performance was compared using the F1 score on the playback dataset across distances. We also assessed the precision of pyannote and WebRTC VAD on the dataset collected in the Bymarka forest following the protocol described in section 2.3.

## 3. Results

### 3.1. Playback experiment results

The ecoVAD model was able to classify human speech with a high degree of confidence up to 10 meters for samples from a man, woman, and child (mean classification confidence > 0.8 for all speakers at 10 meters; Figure 2a). At 20 meters the classification confidence for speech decreased for all speakers (0.727, 0.700, 0.707 means for man, woman, and child respectively) and the speech score of the lower quartile overlapped with the scores for the non-speech samples. However, at 20 meters the speech was barely audible in the audio samples, and the sentences being spoken were not intelligible. The classification confidence for periods of audio with no speech in them remained low at all distances with a mean classification confidence of 0.30.

**Figure 2:**
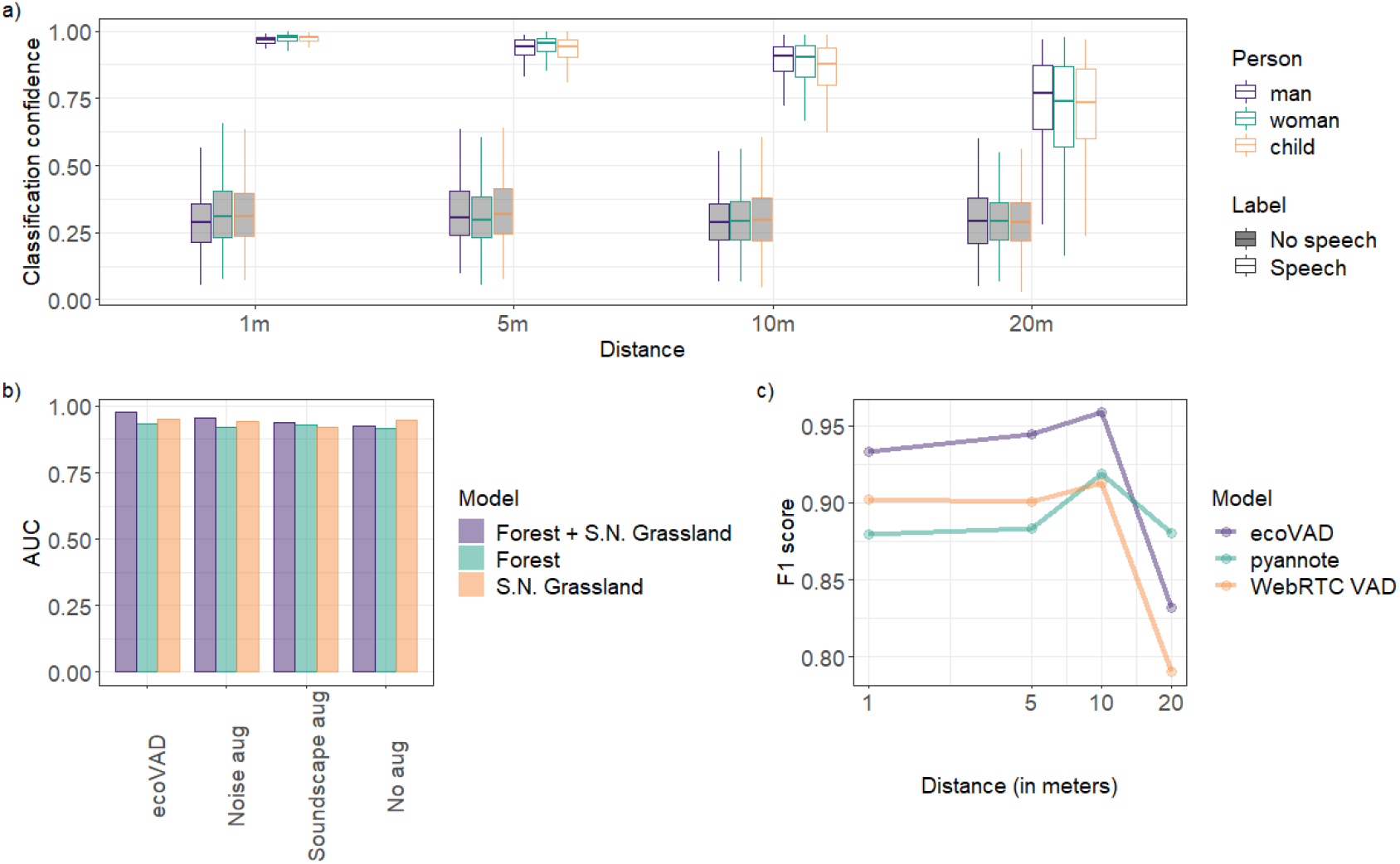
Voice activity detection (VAD) models performed well on eco-acoustic data in controlled playback experiments. **a)** For the ecoVAD model, classification confidence is plotted against distance for samples of speech from man, woman, and child. The model was able to successfully discriminate between 3s audio segments with and without speech at distances of up to 20m (at which point the speech was unintelligible). Performance was consistent across samples spoken by a man, woman, and child. **b)** Accuracy of the models trained using different data augmentation approaches and different soundscape types was consistently high (> 0.9 AUC in all cases). A slight increase in performance was seen for models trained with soundscape data from both forest and semi-natural grasslands (S.N. Grassland) and when both noise and soundscape augmentations techniques were used (ecoVAD). **c)** ecoVAD outperformed two existing state-of-the art VAD models (WebRTC VAD and pyannote) when their performance was evaluated on the same playback experiments.

All models employing some form of data augmentation (noise only, sound only, and ecoVAD) that were trained on both forest and semi-natural grassland reached higher AUCs than the model that was trained with no data augmentation (Figure 2b). We did not notice dramatic differences in performance when testing a model on the landscape it has not been trained on. For instance, the ecoVAD model trained on the forest only data and the model trained on the semi-natural grassland only data had an AUC of 0.97 and 0.98 respectively when tested on forest data and an AUC of 0.91 and 0.93 respectively when tested on semi-natural grassland (Supplementary figure 1).

While ecoVAD was trained on a limited amount of data (i.e., only four days of soundscapes), it outperformed state of the art models such as pyannote and WebRTC VAD on the playback dataset (average F1 score = 0.917, 0.890, 0.876 for ecoVAD, pyannote and WebRTC VAD respectively; Figure 2C). We notice that the performance of all models decreased significantly at 20 meters, particularly for WebRTC VAD (F1 scores at 20 meters = 0.832, 0.88, 0.790 for ecoVAD, pyannote and WebRTC VAD respectively).

### 3.2. Using VAD models for human disturbance quantification

In the passive monitoring dataset collected from Bymarka, Norway, again, data augmentation improved the precision of models. At both locations, the models trained without any data augmentation or with only noise augmentation had high rates of false positives (precision < 0.05, Fig. 3a). Soundscape augmentation, however, resulted in a marked improvement in precision (0.25 in Forest 1, 0.18 in Forest 2). The combination of both noise and soundscape augmentation used in the ecoVAD model resulted in further increased precision in Forest 2 (precision = 0.32), but in Forest 1 resulted in a decrease in precision compared to the model trained with only soundscape augmentation (precision = 0.14).

**Figure 3:**
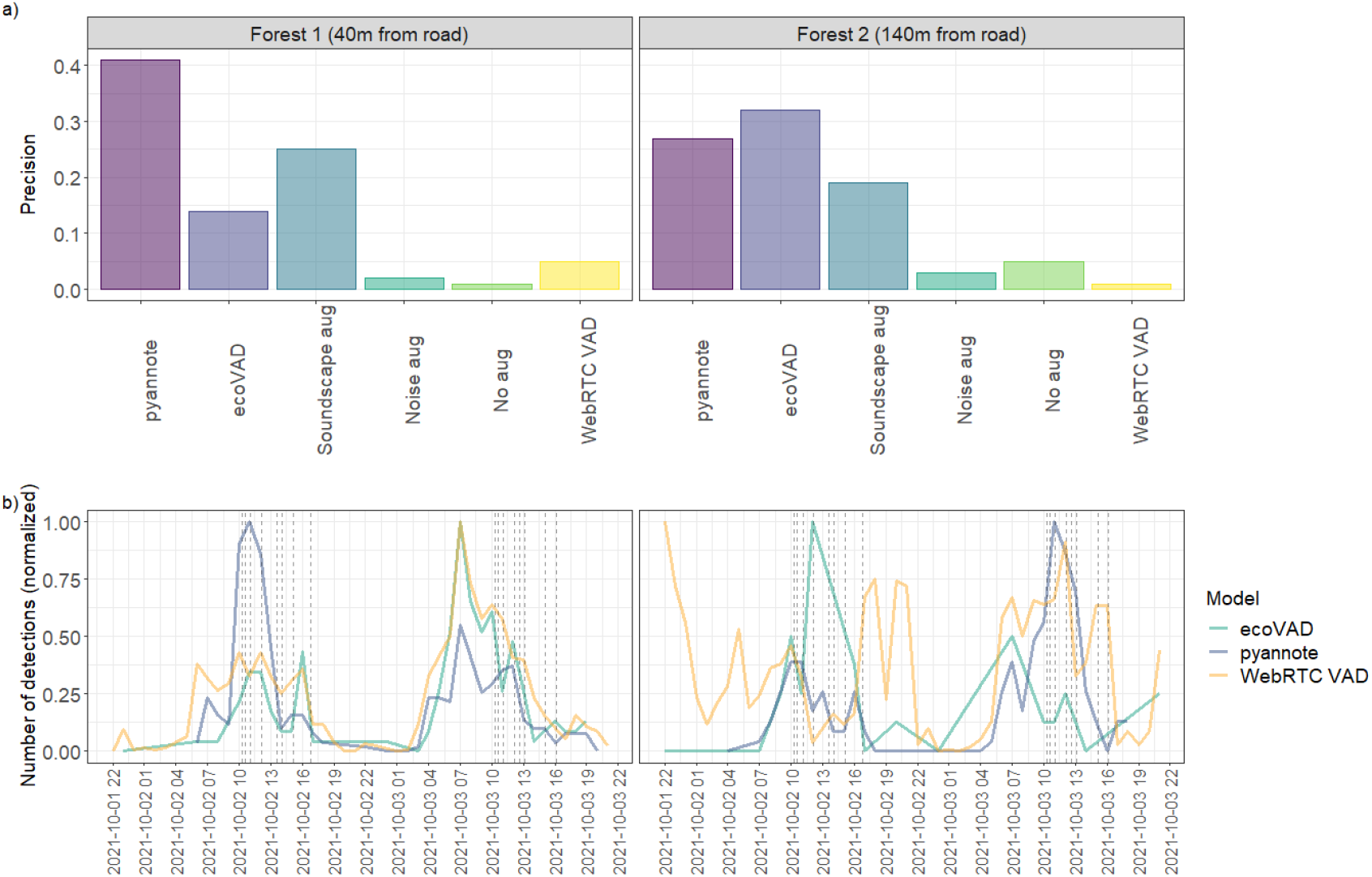
Frequency of voice activity detections serves as an accurate measure of human pressure on a fine temporal resolution. **a)** 100 automated voice detections from two days (October 2^nd^-3^rd^ 2021) of eco-acoustic monitoring were manually labelled to derive the precision of five models at two sites in Bymarka, Norway. Soundscape augmentation had a larger impact upon model precision than noise augmentation. Our model, ecoVAD, achieved state-of-the-art performance, outperforming WebRTC VAD at both locations and pyannote in Forest 2. **b)** Min-max normalised number of hourly detections across the same two-day period for ecoVAD (purple), pyannote (green) and WebRTC VAD (orange) are depicted alongside black dashed lines, which show bus arrival and departure times. There are clear peaks in both voice detections and bus timings during the morning hours on both days, indicating that voice detections can be used as a fine scale measure of human pressure.

In Forest 2, located 140 meters from the nearest road, ecoVAD outperformed both pyannote and WebRTC VAD (increased precision of 0.05 compared to pyannote and 0.31 compared to WebRTC VAD, Fig. 3a). However, pyannote outperformed both ecoVAD (regardless of the augmentation approach used) and WebRTC VAD when evaluated on data from Forest 1 which was 40 meters from the nearest road.

We found a clear link between bus timings and the number of hourly detections using both pyannote and ecoVAD models for both Forest 1 and 2. For WebRTC VAD the link between bus timing and number of hourly detections was clear for Forest 1 but less so for Forest 2, most likely due to the low precision of the model at this location. Between the hours of 06:00 and 18:00 inclusive (approximate daylight hours) there were significantly more speech detections from both the ecoVAD and pyannote models than during the nightime hours (90%, 85% for ecoVAD and 91%, 97% for pyannote on Forest 1 and Forest 2 respectively). Both models registered peaks in normalised detections/hour between 06:00 and 12:00 on both October 2^nd^ and October 3^rd^ at both locations. This is both supported by a higher frequency of bus arrivals in the morning and agrees with the intuition that hiking activity is likely to be higher in the morning than in the late afternoon and evening hours. While most of WebRTC VAD detections were located between 6:00 and 18:00 for Forest 1 (86%) this proportion decreased for Forest 2 (63%).

## 4. Discussion

In this study we showed that VAD models can be successfully used for anonymising eco-acoustic data and for quantifying human disturbance within an ecosystem on a fine temporal resolution. However, our results suggest that not all available models should be considered for the task of anonymization and human disturbance quantification of eco-acoustic data. We demonstrated that augmenting publicly available speech datasets with soundscape recordings and common noise sources (e.g., wind, birdsongs) resulted in speech detection models which outperformed the state-of-the-art. We also found that the peaks in number of voice detections coincided with peak hours of bus activity; suggesting that voice activity detection models can be used to quantify human disturbance within an ecosystem on a fine temporal resolution.

Data augmentation was used in our study when training ecoVAD; a deep learning based voice activity model tailored for use in diverse natural soundscapes. Our playback experiments were conducted in the relatively silent soundscapes of Børsa, Norway, and, intuitively, we found the data augmentation process resulted only in a slight improvement of model accuracy in this setting. However, we found that data augmentation significantly improved ecoVAD’s precision for the passive monitoring data collected from the Bymarka forest. This was most likely due to the more diverse set of confounding sounds such as cars, motorcycle, footsteps, wind, and animals contained within the longer recordings. Our results agree with prior work exploring the use of data augmentation in acoustic machine learning tasks, and suggest that data augmentation may be even more important when working in more acoustically complex environments (e.g., the tropics; 28) where confounding sounds can be even more varied.

Throughout our study we compared the performance of ecoVAD with two existing state-of-the-art models commonly used for voice activity detection; pyannote and WebRTC VAD. In the playback experiments we found that ecoVAD consistently outperformed both WebRTC VAD and pyannote (at all distances except at 20m, at which point the speech was unintelligible to a human listener). However, in the passive monitoring scenario, whilst ecoVAD had significantly higher precision than WebRTC VAD in both locations, pyannote outperformed ecoVAD at the location nearer to the road. Analysing the sources of false detections hints that pyannote was less likely than ecoVAD to falsely identify anthropogenic sounds as speech, but more likely to be confused by natural sounds such as wind or branches falling as speech (Supplementary Figure 2). Even though ecoVAD was trained with some anthropogenic noise sources, the data augmentation was heavily weighted towards biotic sources, whilst the training data used for pyannote (e.g., VoxCeleb; 26) will have contained a greater diversity of anthropogenic background sounds. WebRTC VAD, a classical machine learning algorithm, has been especially trained for voice active detection in real-time (27) and trades accuracy for speed. As shown in our study, WebRTC VAD might not be adapted to large datasets as it returns a large number of false detections and this could hide essential information about the ecosystem of interest. There is unlikely to be a one-size-fits-all approach, however, further work may either employ ensemble approaches or find a better balance in the data augmentation process to enable more robust voice detection across a wide variety of soundscapes.

We found that the frequency of voice detections per hour was linked closely to bus activity at a popular hiking spot in Bymarka, Norway. Our results demonstrate that acoustic data can thus be used as both a direct measure of anthropogenic noise pollution, and an indirect measure of human pressure on an environment – both on fine temporal resolutions. Biodiversity is both directly and indirectly impacted by noise pollution (29) and human land use (30), and understanding these interactions on finer temporal scales could open the door to new opportunities for effective conservation measures. Furthermore, by following our methodology, similar classification models could be developed to detect other anthropogenic sounds sources (e.g., vehicles, building activity) providing a multi-dimensional, continuous, and long-term picture of human pressures on an environment.

## Acknowledgments

We thank Wouter, Raghnild and Marius Koch who volunteered to record the samples for the playback experiments. We also thank Frøde Fossoy who took part in the initial discussions that resulted in this work. Trondheim commune gave permission for recording of audio data in the Bymarka forest (permit reference 21 /40874). This work was supported by the Norwegian Research Council (grant number 160022/F40). SSS was supported by a Herchel Smith Fellowship from the School of Biological Sciences at the University of Cambridge.

## Authors’ contributions

All authors contributed to the conceptualization of the manuscript. CR acquired funding and provided technical support. BC and SS analyzed the data and wrote the original draft of the manuscript. All authors reviewed and edited the manuscript and approved the final version.

## Data Availability Statement

All the code required to reproduce our pipeline can be found at https://gitlab.com/nina-data/eco-acoustic/. For anonymity reasons the data used in this analysis cannot be shared.

